# Statistical Model Comparison Supports a Pitcher Origin of *Utricularia* Suction Traps

**DOI:** 10.64898/2026.04.19.719479

**Authors:** Masafumi Obara, Nicholas J. Matzke, Matt Fulmer, Shane Wright

## Abstract

Carnivorous plants have been the subject of fascination and research ever since Darwin codified the subject in his 1875 book *Insectivorous Plants*. The origin of complex trapping mechanisms from structures adapted for photosynthesis is of particular interest. While Darwin proposed a plausible scenario for the origin of the snap traps of the Venus flytrap from simpler adhesive traps, the origin of the tiny and complex bladder traps of the genus *Utricularia* mystified Darwin and many subsequent workers, despite *Utricularia* being the most diverse genus of carnivorous plants. In this study, we test the “pitcher hypothesis,” which proposes that *Utricularia* bladder traps evolved gradually from an adhesive trap ancestor, via an extinct pitcher trap intermediate. To overcome the lack of any fossil evidence for this scenario, we constructed a variety of continuous-time Markov chain (CTMC) models, each of which consists of a transition matrix allowing or disallowing certain transitions between 11 types of traps. We assembled available phylogenetic trees for 436 carnivorous plant species and noncarnivorous outgroups, classified each species by trap type, and statistically compared the fit of 18 CTMC models using Maximum Likelihood and statistical model comparison with Akaike Information Criterion. The best-fitting model (PH-7R-AAI), consistent with our pitcher hypothesis, had an AIC weight of 60%, with two similar models accounting for the remaining 40%. These results support a circuitous stepwise evolutionary pathway to the bladder trap, and demonstrate how a detailed stepwise evolutionary scenario may be statistically tested even without direct fossil evidence of key intermediate stages.

## Introduction

Dear Darwin […]

The account of *Utricularia* is most marvellous & quite new to me. I am rather surprised that you do not make any remarks on the origin of these extraordinary contrivances for capturing insects.^2^ Did you think they were too obvious? I dare say there is no difficulty, but I feel sure they will be seized on as *inexplicable* by *Nat. Select. &* your silence on the point will be held to show that *you consider them so*! [….] What a wonderfull & long continued series of variations must have led up to the perfect *“trap”* in *Utricularia*, while at any stage of the process the same end might have been gained by a little more development of roots & leaves, as in 9999 plants out of 10,000!

Is this an imaginary difficulty, or do you mean to deal with it in future editions of the “Origin”?

—Alfred R. Wallace letter to Darwin, July 21, 1875, commenting on Darwin’s new book *Insectivorous Plants*

My dear Wallace, […]

If at any time you are curious on subject, you will find development of the Droseraceæ discussed in closing part of Chapt. XV, and I think I have thrown some light on the acquirement of wonderful power of digestion.— With respect to *Utricularia*, I can explain nothing, for there are no gradational genera, and even the embryology or development of the present bladders not made out.

—Darwin reply to Wallace, July 22, 1875

In the 150 years since Darwin’s (1875) *Insectivorous Plants*, carnivorous plants have attracted attention for their extraordinary trapping adaptations, including the snap traps of *Dionaea muscipula* (Venus flytrap) and *Aldrovanda* (the waterwheel plant), the adhesive traps of *Drosera* (sundews) and *Pinguicula* (butterworts), and the pitfall-trap pitchers of *Nepenthes, Cephalotus*, and the Sarraceniaceae. The ability of the Venus flytrap (‘one of the most wonderful in the world’; Darwin, 1875a) to selectively detect insects and rapidly trigger its trap (Forterre et al., 2005) has made the Venus flytrap the iconic carnivorous plant. However, the aquatic bladder traps of the genus *Utricularia* (bladderworts) are arguably even more astounding. Darwin directly observed prey “suddenly” appearing inside bladderwort traps but imagined that the mechanism was passive. It was not until the work of Lloyd (1942) that the bladder trap’s full complexity was detailed, wherein disturbance of a trigger hair causes a double-hinged trap door to open, drawing in prey via negative pressure before the trap resets. Lloyd analogised bladder traps “without exaggeration” to an imaginary self-resetting mousetrap with 20+ necessary components, cooperating with “an astounding degree of mechanical delicacy depending on a fineness of structure scarcely equalled elsewhere in the plant kingdom” (Lloyd, 1942).

### Prior Attempts at Explaining the Origin of the *Utricularia* Trap

While progress has been made in resolving the relationships of *Utricularia* and understanding their diversification to occupy various wet-terrestrial and aquatic habitats (Westermeier et al., 2017), there is still a significant unresolved question about how the unique bladderwort trap evolved, dating back to 1875. Alfred Russel Wallace, the co-discoverer of natural selection and frequent correspondent of Darwin, was always on the lookout for puzzling cases and, upon reading *Insectivorous Plants*, wrote to Darwin asking about the origin of *Utricularia* (Wallace, 1875). Darwin replied by first pointing to his explanations of carnivorous plant traps in the Droseraceae, where he proposed a gradational series from passive sticky leaves (*Drosophyllum*), to slowly-closing sticky leaves (*Drosera*), to the *Dionaea* and *Aldrovanda* snap traps, where “rapid movement makes up for the loss of viscid secretion.” (Darwin 1875). However, addressing Wallace’s question about bladder traps, Darwin was forced to concede, “With respect to *Utricularia*, I can explain nothing, for there are no gradational genera” (Darwin, 1875b). Most later authorities did little better. Lloyd (1942) wrote, “How the highly specialized organs of capture could have evolved seems to defy our present knowledge.” Givnesh (1989) said the relationship between the *Genlisea* eeltrap and the traps of *Pinguicula* and *Utricularia* remained “totally obscure.” Juniper et al. (1989) discuss the evolution of other trap forms in some detail, but of *Utricularia*, they write that it remains “an intractable problem in evolution” and that “there is no complete natural analogue to this trap to our knowledge anywhere else in the plant kingdom, nor any satisfactory evolutionary path” (Juniper et al. 1989). In Taylor’s massive monograph on *Utricularia*, he could only say that the variation in the trap “gives us, or at least me, no inkling as to how this evolved” (Taylor, 1989). We could only find one somewhat detailed hypothesis from before the year 2000, where Snyder (1987) attempts to derive the *Utricularia* bladder from theorized air-sac floats on the roots; however, this can be ruled out on the grounds that the bladders are homologous to leaves (Juniper et al. 1989).

Phylogenetic reconstructions (Fleischmann et al., 2010; Jobson et al., 2017) clearly indicate that *Genlisea* and *Utricularia* are sister genera, both of which share a common evolutionary ancestor with the sticky-leaved *Pinguicula*. Fleischmann’s (2012a) comprehensive review of the genus *Genlisea* made a general proposal for the origin of *Utricularia’s* traps, proposing that the *Genlisea*-*Utricularia* lineage likely evolved from a *Pinguicula*-like ancestor with sticky, glandular leaves. Gradual inward folding of these leaves may have formed tubular, pitcher-like traps, serving as an intermediate stage. In *Genlisea*, these structures specialized into subterranean eel traps with hydrodynamic prey capture, while in *Utricularia*, similar tubular traps evolved into aquatic suction traps with active prey capture mechanisms. This highlights sticky traps as precursors to diverse, complex traps and papers by Fleischmann and colleagues (Fleischmann 2012b; Fleischmann et al. 2018) also link the bladder traps to the other traps in Lentibulariaceae. This hypothesis bears several similarities to our pitcher hypothesis so is discussed below.

### The Pitcher Hypothesis

We propose the pitcher hypothesis, which states that the common ancestor of *Utricularia* and its sister genus *Genlisea* evolved from a group of pitcher plants in the family Lentibulariaceae that is now entirely extinct. This hypothesis entails several transitions between distinct pitcher trap types in the extinct group. While such a scenario risks being dismissed as a speculative “just-so story”, we argue that advances in phylogenetic modelling now allow such hypotheses to be subjected to some substantive, if not definitive, testing, through statistical model comparison.

Although Lloyd (1942) mentioned in passing that bladder traps resemble miniaturised pitchers, to our knowledge the first place the pitcher hypothesis was suggested was a web article by a carnivorous plant enthusiast (Cook, 2001). Matzke (2005) proposed a more detailed version of this hypothesis, which we expand upon here. The main challenge in understanding the origin of the *Utricularia* trap lies in imagining plausible intermediate forms between trap types in *Pinguicula, Genlisea*, and *Utricularia*, especially as these are typically treated as discrete categories (Mithöfer, 2022): flypaper, snap, pitcher, eel, and suction traps. However, recent findings increasingly blur these boundaries. Molecular phylogenies show that species with complex traps (e.g. *Dionaea, Aldrovanda, Utricularia*) are closely related to those with simpler adhesive traps, supporting a trajectory from flypaper to more complex mechanisms (Ellison & Gotelli, 2001). Morphological studies (Clarke, 2001; McPherson, 2009; Roccia et al., 2016) have identified several intermediate forms—for example, *Nepenthes inermis* pitchers function as sticky traps rather than pitfalls, and curled leaves in *Pinguicula lutea* resemble primitive pitchers. Some species even employ hybrid strategies, such as pitcher traps with one-way hairs resembling eel traps (*Sarracenia psittacina, Darlingtonia*) or sticky-light-window combinations (*Nepenthes aristolochioides*). These examples help bridge morphological gaps and make the pitcher hypothesis more plausible.

Our pitcher hypothesis for the origin of the *Utricularia* traps emerges by arranging all trap mechanisms on two axes (Figure 1). One axis is the specialisation of traps for different microenvironments: aerial, ground, amphibious, and aquatic. The second axis is an adhesive-to-pitcher continuum. Trap mechanisms that have been observed in living species and intermediate series that have been postulated in carnivorous plant evolution can then be mapped onto this framework. For example, Darwin’s scenario for the origin of *Aldrovanda*’s aquatic snap trap postulated a path from adhesive traps, through an amphibious *Dionaea*-like stage, to a fully aquatic snap trap. To explain the origin of the *Utricularia* trap, the proposed stages of our pitcher hypothesis are: (a) ancestral flypaper traps, (b) intermediate adhesive/pitcher-like traps, (c) a ground pitcher trap, (d) amphibious eel trap, and finally (e) a fully aquatic suction trap.

**Figure 1.**
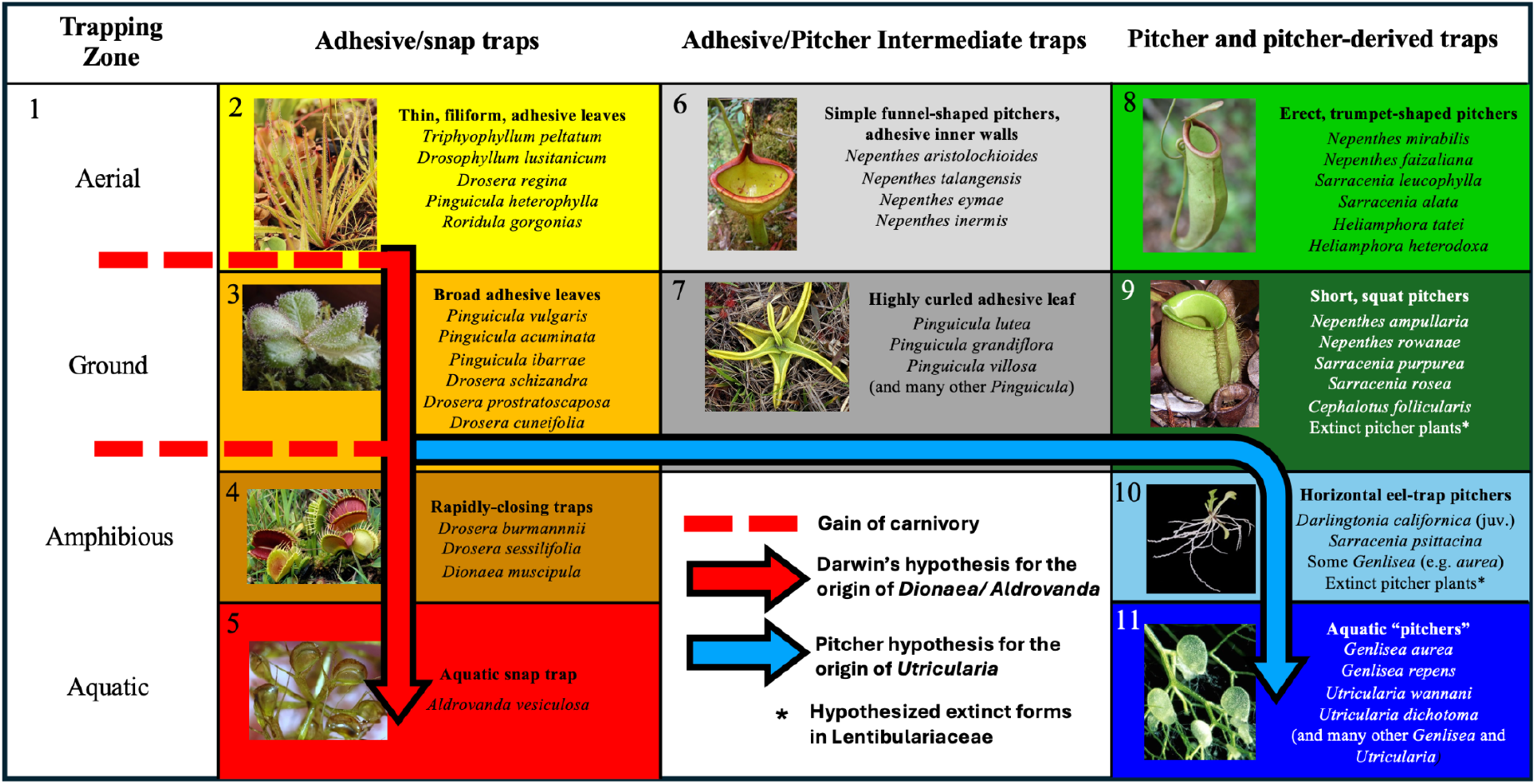
Our pitcher model for the origin of *Utricularia*’s bladder traps (blue arrow) is presented as a series of stages within the overall context of the convergent evolution of carnivorous plant trapping mechanisms, including Darwin’s hypothesis for the origin of snap traps (red arrow). 11 trap states are represented by numbers and colour-coded to match the phylogeny/character mapping figures. Trap character states are as follows: (1) non-carnivorous, (2) aerial adhesive/ flypaper trap, (3) ground adhesive/ flypaper trap, (4) amphibious snap trap, (5) aquatic snap trap, (6) aerial adhesive/pitcher intermediate trap, (7) ground adhesive/pitcher intermediate trap, (8) aerial pitchers, (9) ground pitchers, (10) amphibious pitchers, and (11) aquatic pitchers.

**Figure 2.**
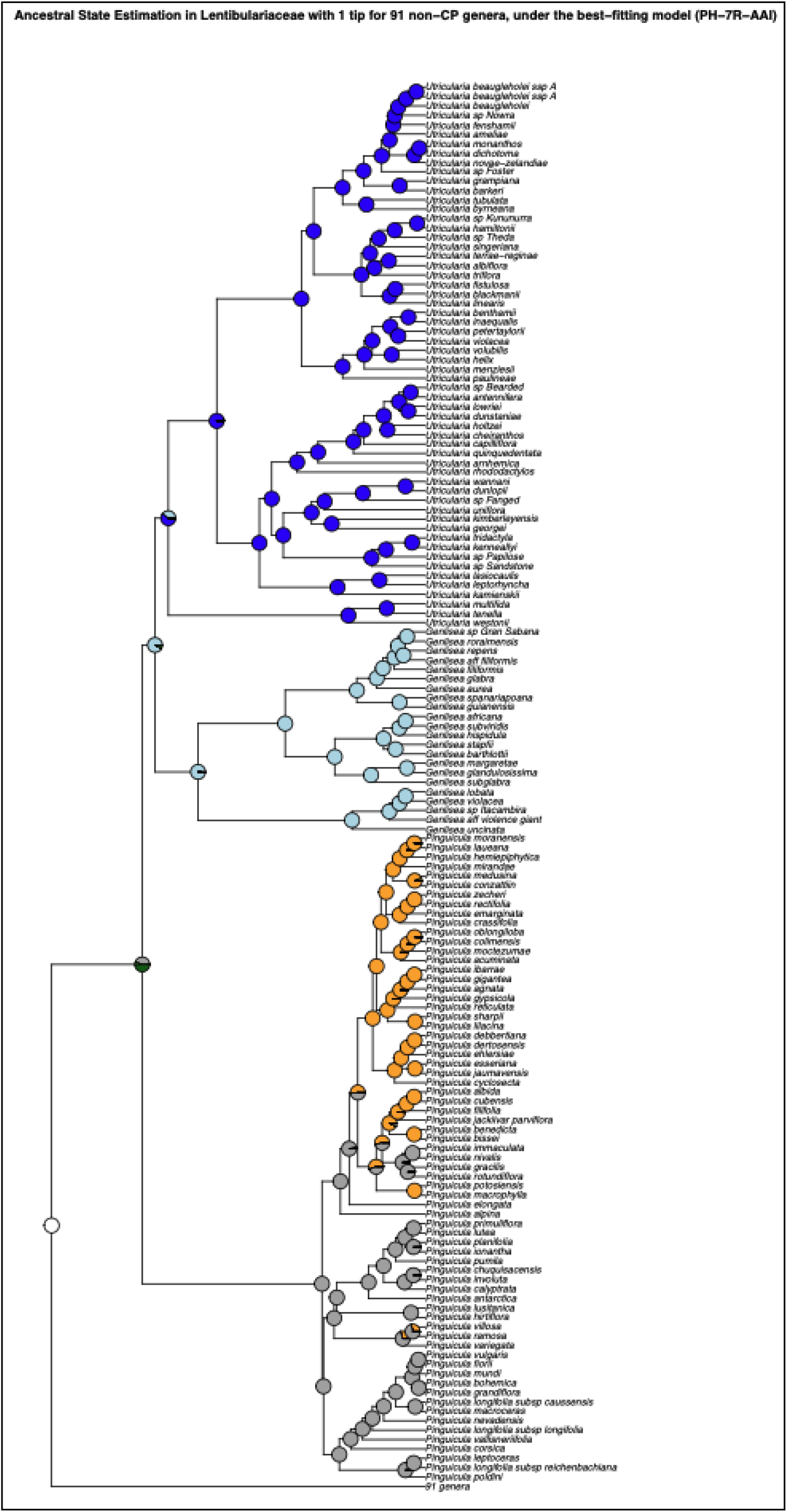
Ancestral trap state estimation for Lentibulariaceae under the best-fitting model (PH-7R-AAI) suggests that the branch below the common ancestor of the clade was non-carnivorous (white) and the common ancestor node had adhesive/pitcher intermediate+ground pitcher traits. The dark blue clade represents *Utricularia* bladder traps, light blue represents *Genlisea* eeltraps, and the clade mixed with orange and grey represents *Pinguicula*

The argument for the plausibility of the transition from (a) to (c) is given above. The argument for (c) to (e) begins with the fact that the traps of *Genlisea* resemble eel traps. The similarities between *Genlisea* and *Utricularia* traps suggest a shared ancestry, with the common ancestor likely possessing a homologous leaf trap (Płachno et al., 2020). Our pitcher hypothesis proposes that this common ancestor was an amphibious eel trap, prostrate on the ground or shallowly buried in moss or soil, much like some living *Genlisea* traps grow horizontally rather than vertically down into the substrate (Lloyd, 1942). We suggest that this postulated ancestor was convergent in form and mechanism to the trap of *S. psittacina*, where the pitcher sits horizontal to the substrate, has a restricted entrance and eel trap mechanism, and can even trap prey while submerged (McPherson & Schnell, 2011). From this common ancestor, the evolution of eel trap in *Genlisea* could have resulted from specialisation, with the twisted arms of the trap possibly evolving from appendages resembling *Darlingtonia* fishtails. Converting the ancestral eel trap into a *Utricularia* suction trap begins with curling the edge of the entrance inwards to form a one-way valve that further impedes escape of live prey and the nutrients diffusing from decaying prey. The addition of suction is then a gradual improvement on eel-trap functionality, helping bring struggling prey into the “pitcher”/digestion chamber. As suction trap capability improves, the reliance on eel trap mechanisms, like hairs that guide prey towards the entrance, can reduce although some living *Utricularia* still use hairs in an apparent eel trap-like mechanism to guide prey to the trap door.

Undoubtedly, this scenario does not address many morphological details, let alone molecular ones, but we suggest that determining the major functional stages by which a complex adaptation evolved is a crucial first step in solving an evolutionary puzzle. The significant advantage of proposing a series of stages and the transitions between them is that this creates an explicit model of trap evolution that can be represented as a transition matrix in a continuous-time Markov chain (CTMC) model, and fit to a dataset consisting of a phylogenetic tree containing the carnivorous plant clades and their non-carnivorous relatives and tip data encoding the trap type of each species. Variant models allowing different transitions and different numbers of parameters can also be implemented to statistically compare the fit of different models for the evolution of carnivorous plant traps. Here, we compare the fit of the pitcher model to other possible models, using the Akaike Information Criterion (AIC) to measure the relative statistical fit of models. The best-fit models are then used in stochastic character mapping to estimate ancestral trap types and the timing and number of transitions between trap types.

## Materials and Methods

### Collecting and Digitising Trees

In order to statistically compare trap evolution models, a dated phylogeny covering all the trapping clades in a single phylogeny, as well as noncarnivorous outgroups, was required. However, carnivorous clades are scattered throughout the angiosperm phylogeny. As our goal is a preliminary test of hypotheses for the evolution of trapping strategies, using a model comparison approach entirely novel in carnivorous plant research, a *de novo* dating study covering thousands of species was deemed unnecessary. Instead we chose to assemble a supertree from previously published studies, in order to obtain a reasonable tree reflecting divergence times from recent publications.

Phylogenetic trees covering each carnivorous plant genus were collected from publications (Ellison et al., 2012; Fleischmann et al., 2010; Jobson et al., 2017; Liu & Smith, 2021; Murphy et al., 2020; Sen et al., 2020; Shimai et al., 2021). These covered the families Droseraceae, Nepenthaceae, Lentibulariaceae, and Sarraceiaceae. Digitisation was accomplished using WebPlotDigitizer (https://automeris.io/WebPlotDigitizer/), and the digitised data for each tree was converted to Newick using “TreeRogue” R scripts (Matzke, 2012). The digitised trees of carnivorous plant clades were grafted onto an angiosperm megaphylogeny from V.PhyloMaker, an R package designed to provide subtrees from a large precalculated phylogenies of vascular plants (Jin & Qian, 2019). When a digitised tree was not dated but had molecular branch lengths, we used r8s (Sanderson, 2004) to produce an ultrametric, approximately time-scaled tree. The digitised *Pinguicula* tree (Shimai et al. 2021) was not dated and had no branch lengths; however, we used the r8s program to impose several time constraints from the dated *Pinguicula* subtree available in V.phyloMaker and combined it with the larger tree.

After assembling the overall supertree (74,533 species), we reduced it to minimize computational effort on the vast regions of the tree that are noncarnivorous. For each carnivorous clade, we kept only three successive non-carnivorous sister groups for each carnivorous clade, and only one species per genus for other non-carnivorous clades. This reduced the tree from 74,533 species to around 1879 species, 432 of which are carnivorous.

This included enough noncarnivorous branch length to record that each carnivorous clade evolved independently from noncarnivorous ancestors.

### Classification of Carnivorous Plant Trap Types

For each sampled carnivorous species we coded, trap type, subtype, trapping zone, maximum trap size, and trap shape for every species based on data acquired from publications (see Supp. Mat.). Trap type refers to sticky leaf, snap, adhesive/pitcher intermediate, pitcher, eel trap, or suction trap. In contrast, the sub-trap refers to the more detailed category, which helps to identify character states. For example, all *Nepenthes* are pitcher plants; however, some *Nepenthes*, like *Nepenthes inermis*, have a sticky inner wall, which suggests an adhesive/pitcher intermediate trap that possesses features of both adhesive and pitcher traps. The trapping zone refers to whether traps are specialised for trapping in aerial, ground, amphibious or aquatic habitats. Based on these classifications, we coded the character states as states numbered 1-11 (see Figure 1).

### Transition Rate Matrices

Once the phylogenetic trees had been assembled, we made transition matrices representing 18 models. The base Pitcher Hypothesis (PH) model (see Table 1) represents our pitcher hypothesis for the origin of the *Utricularia* trap as a series of transitions between 11 states. This model involves 12 transition rate parameters, and this model was compared to alternative models that postulate other allowed transitions. For example, the simple equal-rates (ER) null model allows all trap types to have equal rates of transition to any other trap type (see Table 2). It represents one version of an “anything is possible” model. The rest of the models with rate descriptions can be found in Supp Mat. In the PH model, the loss of carnivory is indicated by rate 1, representing transitions from any other state to state 1. Rate 2 represents the gain of carnivory, postulating that sticky leaf traps were the first form of carnivory to evolve from non-carnivorous ancestors (Darwin, 1875; Craw et al., 1999; Slack, 1988; Juniper et al.,1989). The key transition central to our pitcher hypothesis involves a stepwise evolutionary sequence: from adhesive traps, to adhesive/pitcher intermediates, then to pitcher traps. This might have occurred in either aerial or ground forms of these traps, as representatives of these trap types in both zones are observed in nature (Table 1). In the base PH model, this continuum is represented by four distinct transition rates (rates 9–12 in Table 1), with each transition fixed as irreversible.

**Table 1.** **(a)** A transition matrix representing the Pitcher Hypothesis (PH) model postulated in Figure 1, where the aquatic bladder trap is produced by stepping through states 0-3-7-9-10-11 or 0-2-6-9-10-11. **(b)** Each number in (a) represents a different free transition rate parameter in the PH model.

|  |  |  |  |  |  |  |  |  |  |  |  |
| --- | --- | --- | --- | --- | --- | --- | --- | --- | --- | --- | --- |
| (a) |  |  |  |  |  |  |  |  |  |  |  |
| **Pitcher Hypothesis Model** |  |  |  |  |  |  |  |  |  |  |  |
| States | 1 | 2 | 3 | 4 | 5 | 6 | 7 | 8 | 9 | 10 | 11 |
| 1 | 0 | 2 | 2 |  |  |  |  |  |  |  |  |
| 2 | 1 | 0 | 3 |  |  | 11 |  |  |  |  |  |
| 3 | 1 | 3 | 0 | 4 |  |  | 9 |  |  |  |  |
| 4 | 1 |  |  | 0 | 5 |  |  |  |  |  |  |
| 5 | 1 |  |  |  | 0 |  |  |  |  |  |  |
| 6 | 1 |  |  |  |  | 0 |  | 12 |  |  |  |
| 7 | 1 |  |  |  |  |  | 0 |  | 10 |  |  |
| 8 | 1 |  |  |  |  |  |  | 0 | 6 |  |  |
| 9 | 1 |  |  |  |  |  |  | 6 | 0 | 7 |  |
| 10 | 1 |  |  |  |  |  |  |  |  | 0 | 8 |
| 11 | 1 |  |  |  |  |  |  |  |  |  | 0 |

| (b) |  |
| --- | --- |
| Rates | Description |
| 1 | Loss of Carnivory |
| 2 | Gain of Carnivory |
| 3 | Aerial to Ground (Adhesive) |
| 4 | Ground to Amphibious (Adhesive to Snap) |
| 5 | Amphibious to Aquatic (Snap) |
| 6 | Aerial to Ground (Pitcher) |
| 7 | Ground to Amphibious (Pitcher to Eel) |
| 8 | Amphibious to Aquatic (Eel to *Utricularia*) |
| 9 | Adhesive to Intermediate (Ground) |
| 10 | Intermediate to Pitcher (Ground) |
| 11 | Adhesive to Intermediate (Aerial) |
| 12 | Intermediate to Pitcher (Aerial) |

**Table 2.** Transition matrix for the equal-rates model, the simplest null model. All character states have equal rates to transition into one another, represented by 1.

| Equal-rates Model |  |  |  |  |  |  |  |  |  |  |  |
| --- | --- | --- | --- | --- | --- | --- | --- | --- | --- | --- | --- |
| States | 1 | 2 | 3 | 4 | 5 | 6 | 7 | 8 | 9 | 10 | 11 |
| 1 | 0 | 1 | 1 | 1 | 1 | 1 | 1 | 1 | 1 | 1 | 1 |
| 2 | 1 | 0 | 1 | 1 | 1 | 1 | 1 | 1 | 1 | 1 | 1 |
| 3 | 1 | 1 | 0 | 1 | 1 | 1 | 1 | 1 | 1 | 1 | 1 |
| 4 | 1 | 1 | 1 | 0 | 1 | 1 | 1 | 1 | 1 | 1 | 1 |
| 5 | 1 | 1 | 1 | 1 | 0 | 1 | 1 | 1 | 1 | 1 | 1 |
| 6 | 1 | 1 | 1 | 1 | 1 | 0 | 1 | 1 | 1 | 1 | 1 |
| 7 | 1 | 1 | 1 | 1 | 1 | 1 | 0 | 1 | 1 | 1 | 1 |
| 8 | 1 | 1 | 1 | 1 | 1 | 1 | 1 | 0 | 1 | 1 | 1 |
| 9 | 1 | 1 | 1 | 1 | 1 | 1 | 1 | 1 | 0 | 1 | 1 |
| 10 | 1 | 1 | 1 | 1 | 1 | 1 | 1 | 1 | 1 | 0 | 1 |
| 11 | 1 | 1 | 1 | 1 | 1 | 1 | 1 | 1 | 1 | 1 | 0 |

Variant models modify this structure to test alternative evolutionary scenarios. For example, the Pitcher Hypothesis Reversible (PHR) model allows the adhesive–intermediate–pitcher transitions to be reversible, while the PH-8R model also permits reversibility but distinguishes eight separate rates across these transitions. The PH-7R models (four variants) similarly incorporate reversibility but reduce the number of distinct rates to seven, with one transition within the continuum fixed as irreversible. Finally, the PHJ model serves as a null model to test whether *Utricularia*’s bladder traps could have evolved without passing through the adhesive/pitcher intermediate stage, representing a direct transition from adhesive to pitcher traps. Variant models and additional null models are described in the Supplementary Data (11state_rate_matrix).

### Evaluating Markov Models for Ancestral Character Estimation

We used the “fitMk.parallel” function in phytools (Revell, 2024) to infer maximum likelihood parameters for various evolutionary models of trap-type transitions. Model fit was evaluated using the maximised log-likelihood (lnL) and Akaike Information Criterion (AIC) scores (Lanfear et al., 2014). AIC provides a framework for comparing alternative evolutionary models based on their relative explanatory power and predictive accuracy (Burnham & Anderson, 2002). Unlike the Bayesian Information Criterion (BIC), which assumes the true model is among those tested and applies a stronger penalty for complexity (Burnham & Anderson, 2002), AIC is more appropriate for phylogenetic comparative analyses where models represent biological hypotheses rather than exact descriptions of reality. This approach allows biologically plausible but complex models, such as variants of our pitcher hypothesis, to be evaluated without over-penalisation for additional parameters.

After selection of the best-fitting model, the probabilities of ancestral trap states were calculated using ancestral character estimation. Transition dynamics were further explored with stochastic character mapping using the function “simmap” in phytools. 100 stochastic sampling simulations were performed. For key branches where major transitions in trap types occurred, state frequencies across maps were aggregated at 100 uniformly-spaced points along the branch, in order to visualise probability of each state over time.

## Results

### Model Selection

Maximised log-likelihood (LnL) and AIC with AIC weights for each of the 18 models are shown in Table 3. The best-fitting model was PH-7R-AAI, which is one of the pitcher hypothesis model variants. This model was selected based on the lowest AIC value of 707.10235 and the highest AIC weight of nearly 60%. Two similar models (PH-8R and PH-7R-GIP) account for the remaining ~40% of the weight, bringing the combined support for the top three models to >99.9% of the credible set. These results indicate that models allowing the transitional pathways represented in our pitcher hypothesis for the origin of *Utricularia* strongly outperform many other possible models, including those that permit a broader range of transitions between traps, such as the parameter-poor Equal Rates (ER) model.

**Table 3.** AIC summary table for each phylogenetic model. Provided are model log-likelihoods, delta AIC (difference from the best, lowest AIC), relative likelihood (rel_likes), AIC values and weights.

| Model | Model Description | Log-likelihood | # of Parameters | AIC | deltaAIC | AIC_wt (%) |
| --- | --- | --- | --- | --- | --- | --- |
| PH-7R-AAI | Pitcher Hypothesis: 7 rates Aerial one-way Adhesive to Intermediate | -338.6 | 15 | 707.2 | 0 | 60% |
| PH-8R | Pitcher Hypothesis: 8 Rates | -338.6 | 16 | 709.2 | 2 | 22% |
| PH-7R-GIP | Pitcher Hypothesis: 7 Rates Ground one-way Intermediate to Pitcher | -339.8 | 15 | 709.6 | 2.4 | 18% |
| PH-6R-A | Pitcher Hypothesis: 6 Rates Aerial one-way | -397.5 | 14 | 823 | 115.8 | 0% |
| PH-7R-AIP | Pitcher Hypothesis: 7 Rates Aerial one-way Intermediate to Pitcher | -397.5 | 15 | 825 | 117.8 | 0% |
| PH-6R-G | Pitcher Hypothesis: 6 Rates Ground one-way | -370.2 | 14 | 768.4 | 61.2 | 0% |
| PH-7R-GAI | Pitcher Hypothesis: 7 Rates Ground one-way Adhesive to Intermediate | -370.2 | 15 | 770.4 | 63.2 | 0% |
| PH | Pitcher Hypothesis (base) | -382.8 | 12 | 789.6 | 82.4 | 0% |
| ARD | All Rates Different | -311.2 | 110 | 842.4 | 135.2 | 0% |
| SYM | Symmetric | -388.5 | 55 | 887 | 179.8 | 0% |
| PHR | Pitcher Hypothesis Reversible | -425.2 | 12 | 874.4 | 167.2 | 0% |
| ARVTZ | Assymetric Rate Variation by Trapping Zone | -468.7 | 14 | 965.4 | 258.2 | 0% |
| SRVTZ | Symmetric Rate Variation by Trapping Zone | -471.1 | 14 | 970.2 | 263 | 0% |
| GLCU | Gain_Loss-Change-Unconstrained | -501.2 | 3 | 1008.4 | 301.2 | 0% |
| GLCC | Gain-Loss-Change-Constrained | -602.7 | 3 | 1211.4 | 504.2 | 0% |
| ER | Equal Rates | -662 | 1 | 1326 | 618.8 | 0% |
| GLCTZ | Gain-Loss-Change-within Trapping Zone | -865.1 | 3 | 1736.2 | 1029 | 0% |
| PHJ | Pitcher Hypothesis Jump | -1045.4 | 10 | 2110.8 | 1403.6 | 0% |

### Lentibulariaceae

Ancestral state probabilities and stochastic mapping under the best-fitting PH-7R-AAI model suggest that the branch below the common ancestor of Lentibulariaceae was non-carnivorous (indicated by the white circle at the branch bottom in Figure 2). The most recent common ancestor of crown Lentibulariaceae was likely carnivorous with either an “intermediate” trap type, having both adhesive and pitcher traits (indicated by the grey in Figure 2), or a ground pitcher trap (dark green) (see Figure 2). The ancestral trap then diverged into more specialised traps over time (*Utricularia, Genlisea*, and *Pinguicula*). State distribution plots visualise the change in ancestral state probabilities along the root branch of Lentibulariaceae, suggesting that a non-carnivorous ancestor (very light grey line, equivalent to white circles) likely acquired adhesive traits (orange + yellow line) in the middle of the branch, which become the most probable state early in the branch. The ground adhesive/pitcher intermediate traits (dark grey line) then become the most probable state (see Figure 3a) and finally, the ground pitcher traits become the most probable state just before the common ancestor of Lentibulariaceae.

**Figure 3.**
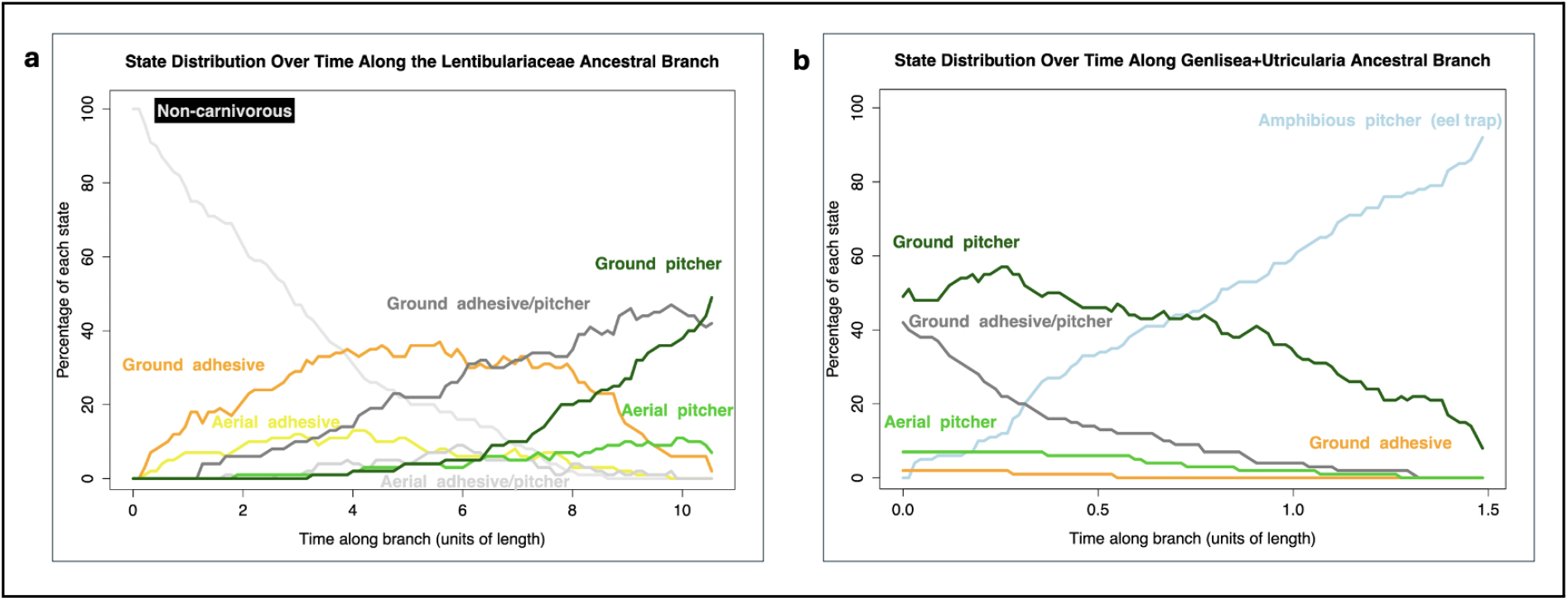
**(a)** State probabilities over time along the Lentibulariaceae ancestral branch, showing the percentage of each state at different points along the branch’s length. The plot illustrates changes in the prevalence of each state over time, with the ground adhesive/pitcher intermediate and ground pitcher traits peaking in frequency. **(b)** State probabilities over time along the *Genlisea+Utricularia* ancestral branch, showing the percentage of each state at different points along the branch’s length. The plot illustrates changes in the prevalence of each state over time, with the amphibious eel trap peaking in frequency, with ground pitcher traits decreasing over time.

Many of the early branches in the *Pinguicula* radiation reconstruct as ground adhesive/pitcher intermediate traps, like some living *Pinguicula*. On the branch ancestral to the common ancestor of *Genlisea*+*Utricularia*, the ground pitcher traits (dark green line) begins as the most probable state, and increases briefly, but then decreases the hypothesised amphibious eeltrap pitcher increases in probability, leading to the last common ancestor node for the two genera (see Figure 3b).

## Discussion

Our analyses support the first statistical test of the Pitcher Hypothesis, which proposes a gradual evolutionary transition from adhesive traps to *Utricularia* bladder traps via a series of trap types, several of which are not observed in living Lentibulariacea, but which have convergently evolved repeatedly in other carnivorous plant clades. Models based on this hypothesis were the top 8 best-fitting among the 18 tested. These models (e.g., PH, PHR, PH-6R-A, PH-7R-AAI, PH-8R) describe a pathway where ancestral flypaper traps evolve into traps with both adhesive and pitcher traits, then into pitcher traps, amphibious eel traps, and finally aquatic bladder traps. The PT variants explore scenarios such as whether transitions are reversible. The best-supported model, PH-7R-AAI, does not permit reversal from the adhesive/pitcher intermediate to adhesive traps, suggesting an evolutionary constraint—intermediate traps may evolve toward pitcher traits but not back to adhesive forms.

The consistent selection of Pitcher Hypothesis models indicates they fit the data (phylogeny and observed trap types) better than alternatives. The 9th and 10th ranked models were SYM (symmetric transitions) and ARD (all rates different). While ARD had a higher log-likelihood (−311.2 vs. −338.6 for PH-7R-AAI), it was heavily penalised for its 110 free parameters, compared to just 15 in the best-fitting model. AIC penalises complexity to avoid overfitting, which explains the preference for simpler, biologically grounded models like PH-7R-AAI. Zone-based models such as ARVTZ and SRVTZ, which apply rate variation by trapping zone (e.g. aerial, ground), ranked 12th and 13th. Gain-loss-change models (GLCU, GLCC, GLCTZ) were ranked 14th to 17th. These models test whether carnivory can be gained, lost, or altered within trap types or zones. The constrained versions restrict transitions to within categories (e.g., between trap types), but not across zones. Their lower performance suggests that such restrictions oversimplify trap evolution. In essence, their poor fit tests and rejects the hypothesis that traps evolve strictly within ecological zones or functional categories.

These findings underscore that a comprehensive understanding of carnivorous trap evolution must account for both ecological context (trapping zones) and functional morphology (trap types). Although zone-based models (e.g., ARVTZ) capture important ecological aspects, pitcher hypothesis models provide a better overall fit, reinforcing the importance of transitional trap forms in evolutionary history. The equal-rate model (ER), which assumes uniform transition rates, ranked 16th. The poorest-performing model, PHJ, does not permit the adhesive/pitcher intermediate trap as part of transitions to bladder traps (e.g., adhesive to pitcher), performing poorly and further emphasising the importance of considering intermediate adhesive/pitcher forms in trap evolution.

## Supporting information

11state_rate_matrix

## Limitations and Future Directions

This study on the evolution of carnivorous plant traps has several limitations. A primary constraint lies in the available data. Our analysis relies on existing data from publications across carnivorous plant lineages, which is incomplete. Expanding genomic and morphological data, particularly for Lentibulariaceae, would improve resolution and reduce potential biases.

We view our models as stepping stones to explain the evolution of carnivorous plant traps. We suggest that the phylogenetic framework of statistical model comparison is valuable because it allows explicit evaluation of model fit and provides a quantitative measure of the degree to which the data support different models. For tractability, trap evolution was discretised into 11 trap types. While this enables hypothesis testing about broad evolutionary stages, it oversimplifies variation. Future work could incorporate continuous traits (e.g. trap size) or break down traps into component traits (e.g. trichomes, mucilage).

A further limitation relates to phylogenetic dating. Our dated supertree was constructed by merging existing dated clades with undated ones scaled using r8s. While practical, this method introduces temporal uncertainty—especially in regions like the short branch between *Genlisea*+*Utricularia* and the Lentibulariaceae ancestor. This unexpectedly short timeframe (under 2 million years) for major trap transitions may reflect artefacts of tree construction. Although model comparisons are valid because all were run on the same tree, a future study could attempt a full BEAST (Bouckaert et al., 2019) dating analysis of Lentibulariaceae, though this is hindered by limited fossil calibrations.

## Acknowledgements

MO, SW and NJM were supported by New Zealand Royal Society RDF 21-UOA-040. MO and NJM were additionally supported by HFSP grant RGY0072/21. NJM was also supported by ARC DP240100462, Marsden Grant 18-UOA-034, University of Auckland DRDF Research Fund #3727963, and University of Auckland, Faculty of Science Research Development Fund, FoS RDF #3732317. MF was supported by Royal Society Marsden Fast Start MFP-UOA2211.

We are grateful to Tanya Renner for her critical review of the manuscript during its preparation.

